# TCR*β* rearrangements without D-segment are common, abundant and public

**DOI:** 10.1101/2021.03.05.434088

**Authors:** Peter C. de Greef, Rob J. de Boer

## Abstract

T cells play an important role in adaptive immunity. An enormous clonal diversity of T-cells with a different specificity, encoded by the T-cell receptor (TCR), protect the body against infection. Most TCR*β* chains are generated from a V-, D-, and J-segment during recombination in the thymus. Although complete absence of the D-segment is not easily detectable from sequencing data, we find convincing evidence for a substantial proportion of TCR*β* rearrangements lacking a D-segment. Additionally, sequences without a D-segment are more likely to be abundant within individuals and/or shared between individuals. We find that such sequences are preferentially generated during fetal development and persist within the elderly. Summarizing, TCR*β* rearrangements without a D-segment are not uncommon, and tend to allow for TCR*β* chains with a high abundance in the naive repertoire.

## 1 Introduction

The adaptive immune system relies on large and diverse repertoires of B- and T-lymphocytes. When encountering antigen, specific lymphocytes start proliferating to clear the pathogen. Many of the cells die after clearance, but others are maintained and form a memory that can be recalled after repeated antigen exposure. The specificity of *αβ* T-cells is determined by the *α* and *β* chain of the T-cell receptor (TCR). These are generated by recombination of variable (V), diversity (D) and joining (J) regions for the TCR*β* and V and J for the TCR*α* chain. During V(D)J-recombination in the thymus, one variant of each of these segments is recombined in a semi-random manner, with deletions and non-templated insertions occurring at the junction(s). The combination of the generated *β* and *α* chains of the TCR yield an enormous potential diversity (> 10^20^ [25, 12]), of which only a small subset is realized in the actual TCR repertoire with a diversity estimated to be around 10^8^ [15].

The recombination process is guided by recombination signal sequences (RSSs) flanking the V, D and J segments. The RSSs contain spacers of 12 or 23 basepairs (bp), and two gene segments can only be recombined when they have different spacer lengths, a principle that is known as the 12/23 rule. In the TCRB locus, the 3’ ends of V and D segments have 23-bp spacer RSSs, while the 5’ end of D and J segments have 12-bp spacer RSSs. Following the 12/23 rule, it is therefore possible to have direct V-to-J rearrangements, not including a D-segment. Ma *et al.* studied TCR*β* sequencing data in which no D-segment could be identified and observed that this occurs in about 0.7% of rearrangements in humans [9]. Previous studies in human cell lines and mice reported V-to-J rearrangements to be rare due to the so called B12/23 restriction [1, 22]. Another scenario that would lead to the complete absence of the D-segment is a large number of deletions, which may happen before and/or after Terminal deoxynucleotidyl transferase (TdT)-mediated N-additions. It is not possible to infer the underlying mechanism from TCR sequencing data as different recombination scenarios lead to identical TCR*β* rearrangements [23]. Moreover, the measured fraction of V-J rearrangements also critically depends on the method used for estimating which nucleotides are derived from the D-segment.

When sequencing TCR*α* or *β* chains from samples of T cells, large differences in abundance are observed, even within samples of naive T cells [16, 23, 15, 14]. It should be noted that this provides no evidence for the existence of large *αβ* clones in the naive compartment. TCR chains differ orders of magnitude in their likelihood to be generated and one common *α* chain is expected to pair with many different *β* chains, or vice versa. The generation probabilities can be estimated using generative models [13, 11, 18] and correlate well with TCR*α* abundance in the naive repertoire [4]. This indicates that abundant single TCR chains can largely be explained by repeated thymic production, and that these occur frequently by summation over different *αβ* clones. Taking this into account, mathematical modelling provided evidence for the existence of large clones in the naive compartment, that are not explained by generation probability [4]. These clones are large for another reason, e.g. high division rates in the periphery [6], which may be due to TCR interactions with self-peptide MHC complexes.

Here we study characteristics of TCR*β* sequences that are abundant in the naive T-cell compartment. We find that TCR*β* rearrangements without a D-segment are a likely outcome of V(D)J-recombination, but not easily identified. TCR*β* chains that are abundant among naive T cells are strongly enriched for having no D-segment. We performed a meta-analysis of TCR*β* sequence data, providing evidence for fetal origin of many of such sequences, which may explain why they are shared between so many individuals. Together, this shows that absence of a D-segment is not uncommon in TCR*β* rearrangements and that it is an import factor explaining TCR*β* abundance in the naive repertoire.

## 2 Results

### 2.1 The naive T-cell repertoire contains abundant TCR sequences that lack glycine in their CDR3

The naive T-cell repertoire consists of a huge clonal diversity, of which just a small fraction can be observed in a typical sample of cells. In addition, when RNA is used to sequence the TCR*β* chains in T cells, differential TCR expression levels may overestimate the measured abundance of a given T cell clone [4]. We therefore re-analyze the data from Qi *et al.* [15], who sequenced the TCR*β* repertoires of memory and naive T-cells from young and aged healthy individuals using five replicates per subset. Measuring the number of samples a given TCR*β* appears in, i.e. the incidence, classifies the abundance of sequences without biases due to multiple RNA contributions by single cells. We processed the subsamples independently using RTCR [7], which performs clustering of likely erroneous sequences using sample-specific estimates of error rates, while maintaining as much as possible of the diversity.

In line with the results presented in Qi *et al.*, we find that the vast majority of the sequences in the naive T-cell samples appears only in a single subsample, underlining the enormous diversity of the naive repertoire. However, there is also a substantial proportion of TCR*β* sequences that are found in two or more subsamples of naive T-cells (Fig. 1A). The median fraction of sequences with an incidence > 1 was 8.0 times higher in aged than in young individuals, confirming the earlier finding that naive T-cell diversity is lost with age [15]. We reasoned that some, and in particular the more abundant sequences may be derived from missorted memory T-cell clones. Therefore, we also analyzed the effect of discarding all sequences that were also observed in at least one of the corresponding memory samples. Although this correction did remove a larger fraction of the abundant sequences than of those with incidence 1, the incidence of most abundant rearrangements remained unchanged (Fig. 1A). This confirms that the naive T-cell receptor repertoire of both young and aged individuals contains abundant TCR*β* sequences [4].

**Figure 1:**
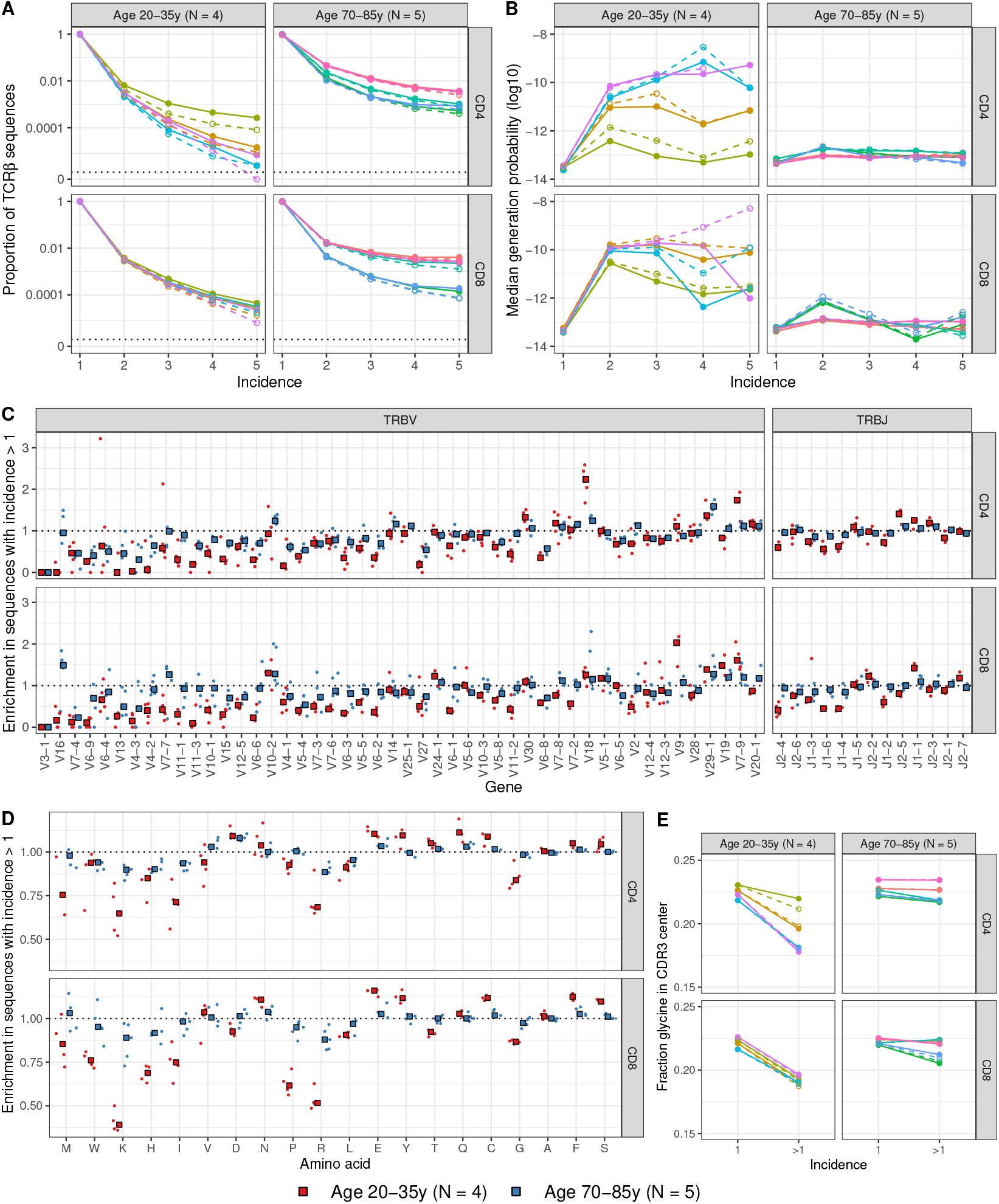
Features of abundant sequences in the naive T-cell repertoire of young and aged individuals. **A.** Fraction of TCR*β* sequences occurring in one or multiple subsamples (i.e., incidence). The vertical axis is log-scaled with 0 added at the bottom. The solid lines and closed circles represent normal data, the dashed lines with open circles show results after removing sequences that were also present in the corresponding sample(s) of memory T cells. **B.** The median generation probability as a function of the incidence. Colors as in A. **C.** The fraction of sequences having a V or J segment with incidence > 1 divided by the fraction of sequences with incidence 1. The gene segments are sorted by increasing median usage in sequences with incidence 1. Rectangles represent median values, individual samples are represented by circles. The horizontal dotted line represents an enrichment of 1, which means that there is no difference between incidence 1 and abundant sequences. **D.** The enrichment of mean amino acid frequencies like in C. The amino acid frequency of each CDR3 is calculated by dividing the CDR3 amino acid counts divided by its total length, to account for differential CDR3 lengths. Amino acids are sorted by increasing median usage among sequences with incidence 1. **E.** The mean proportion of glycine residues among the center 5 amino acids of the CDR3 from TCR*β* sequences with incidence 1 and higher.

TCR*β* sequences differ several orders of magnitude in their probability of being generated during VDJ-recombination. To investigate if this explains the observed differences in abundance in the naive T-cell repertoire, we estimated generation probabilities of the sequences using OLGA [18]. The median generation probability of infrequent TCR*β* sequences (observed in a single subsample) were very similar among all individuals. The abundant TCR*β* sequences, however, were enriched for having a high generation probability in young individuals, albeit to a different extent (Fig. 1B). This indicates that the likelihood of TCR*β* generation, indicating repeated thymus production, is a key factor determining the abundance of TCR*β* sequences in the naive repertoire of young adults. Samples from aged individuals, that have much lower [24] or even no thymus T-cell production [21], contained more abundant sequences, that showed a much smaller enrichment of high generation probability (Fig. 1B). These results remained qualitatively similar after cleaning potential contamination by removing sequences overlapping with the memory compartment (dashed lines in Fig. 1B). Together, these results indicates that differential thymus production affects TCR*β* abundance in young individuals, and that this effect dilutes with age.

While generation probabilities may influence thymus production rates, TCR specificity is expected to affect clonal fitness in the periphery, e.g. through stimulation by self-peptide MHC. We therefore compared the frequencies at which V and J gene segments are used (V/J usage) of abundant and non-abundant TCR*β* sequences in the naive repertoire (Fig. 1C) and observed two overall trends in our data. First, in all individuals we observed a positive correlation between the overall usage of a V or J segment and the enrichment of that segment in abundant TCR*β* sequences compared to rare TCR*β* sequences (Fig. S1A&B). In other words, the V segments that are overall more common are even more strongly over-represented in sequences shared between subsamples. Second, enrichment or depletion of V and J segments in abundant sequences was similar between most individuals, but the effects were stronger for young individuals in most cases. The most characteristic V-segments of abundant TCR*β* sequences were TRBV18 for CD4^+^ T cells, TRBV9 for CD8^+^ T cells and TRBV7-9 for both.

The observation that common V and J segments are even more common among abundant TCR*β* sequences implies some sort of positive feedback mechanism. This surprising result can be intuitively understood when taking summation over multiple TCR*αβ* clones into account [4]. Rearrangements using common V and J segments are a more likely outcome of V(D)J-recombination, i.e., tend to have a high generation probability. Such TCR*β* chains are more likely to be produced repeatedly, and are expected to pair with different TCR*α* chains, leading to multiple TCR*αβ* clones sharing the TCR*β* chain. Thus, observing a particular TCR*β* in several samples means that this particular recombination was realized several times, which implies a higher order dependence on its generation probability. This higher order explains the positive feed-back observed in Fig. 1C, i.e., the additional enrichment of common V and J segments in TCR*β* sequences with a high incidence. Importantly, this implies that the enrichment observed in Fig. 1C and Fig. S1A&B is a consequence of the abundance, and not a mechanism explaining TCR*β* sequence abundance.

We obtained similar results for amino acid usage in the translated CDR3 sequences, which are the main determinant for TCR specificity. The relative amino acid enrichments in abundant TCR*β* sequences were surprisingly similar for the commonly used amino acids within the age groups, but quite dissimilar between young and aged individuals: most enrichment values were closer to 1 (e.g., no difference) in the aged individuals. Like for V/J usage, we noted a positive correlation between overall amino acid usage and enrichment among abundant sequences (Fig. 1D, Fig. S1C). This trend, however, did not apply to glycine. Although being the fourth most common amino acid in the CDR3, it appeared substantially less within abundant sequences from young individuals. In absolute sense (i.e., the frequency of glycine in sequences with incidence 1 minus the frequency of glycine in sequences with incidence > 1), the depletion was even the largest effect size in our dataset. Consequently, the five amino acids in the middle of the CDR3, which are most likely to contact the peptide epitope, contained significantly fewer glycine residues in abundant sequences from young individuals (*p* < 0.01, Wilcoxon signed-rank test; Fig. 1E). This shift was less pronounced in the repertoires of aged individuals, indicating an effect of glycine on abundance that dilutes with age. Since glycine breaks the trend expected from summation, it could play a mechanistic role in explaining TCR*β* abundance.

### 2.2 Many abundant TCR*β* sequences are V-J rearrangements without D-segment

The observation that abundant TCR*β* sequences in the naive repertoire of young individuals contain fewer glycine residues raises the question how these sequences are generated. When translating the nucleotide sequence of both D-segment genes, there is a striking number of guanine-rich codons that translate to glycine (Fig. 2A). This means that deletions at both ends of the D remove guanine nucleotides, and thus likely glycine amino acids from the CDR3 sequence.

**Figure 2:**
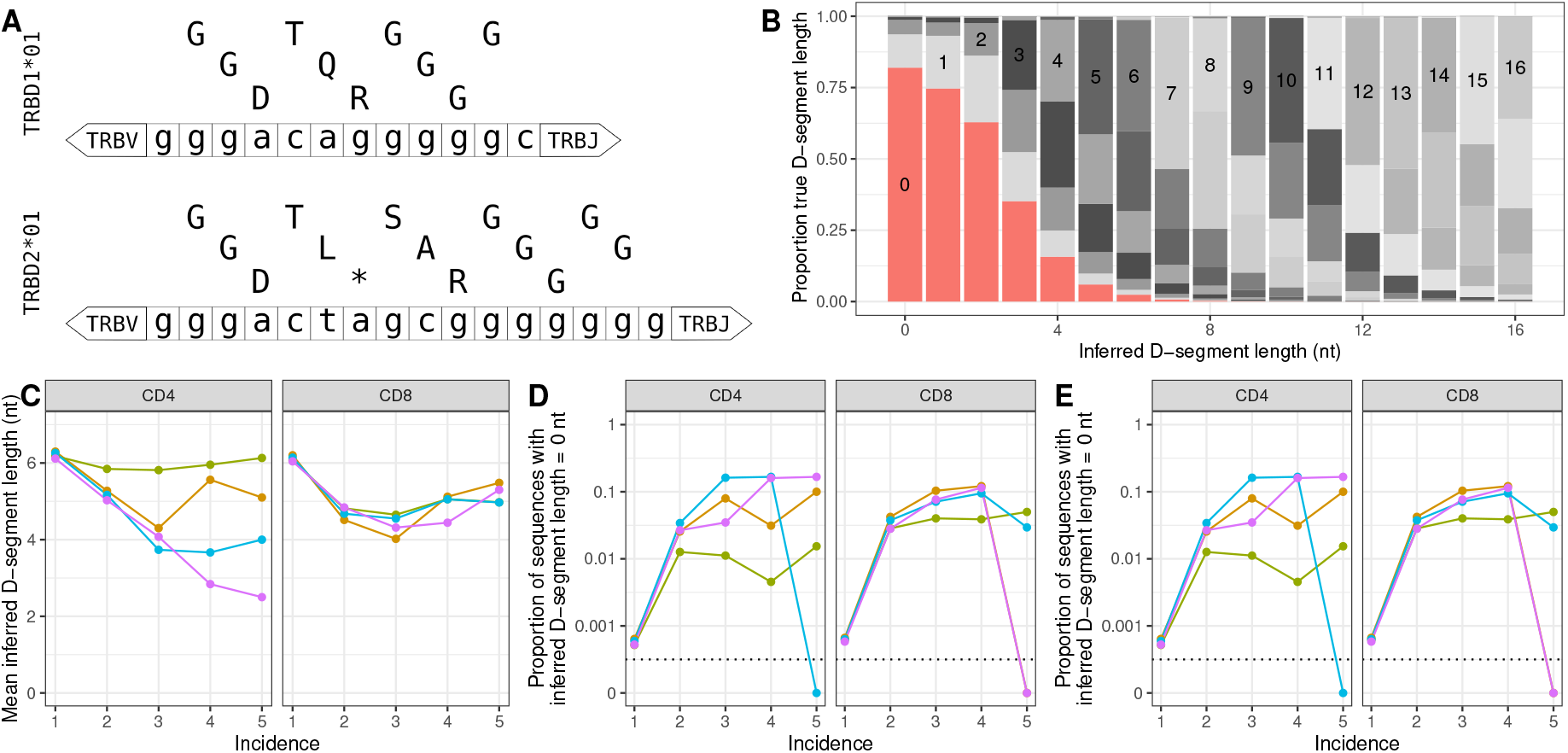
CDR3s without glycine are often indicative of rearrangements lacking a D-segment. **A.** Nucleotide sequences with amino acid translation in all t1hree frames of both human TRBD genes. The asterisk in the sequence represents a stop codon. Note that IMGT [8] lists a second allele for TRBD2, in which the fourth G from the 5’ end is replaced by an A. This is not shown in the figure, but taken into account in the analysis (see Methods). **B.** Comparison between inferred and true lengths of D-segments in an *in silico* repertoire of 10^6^ productive rearrangements generated using IGoR [10]. The proportion of true D-segment lengths is plotted as a function of the inferred D-segment length, which is the maximum region in the non-V/J encoded part of the CDR3 nucleotide sequence matching any D-allele. The bar graph segments are colored by true D-segment length, with inserted numbers indicating identical true and inferred values. **C.** Mean inferred D-segment length as a function of incidence in the naive repertoires of young individuals (colors matching Fig. 1). **D.** Fraction of sequences with an inferred D-segment length of 0 nucleotides as a function of incidence. The vertical axis has a logarithmic scale with 0 added at the bottom of the axis. **E.** Fraction of sequences with an inferred D-segment length of 2 or less nucleotides, most of which likely representing rearrangements without a D-segment.

We therefore wondered if abundant TCR*β* rearrangements in young individuals indeed contain fewer nucleotides that are encoded by the D-segment. It is not straightforward to analyze the D-segment length, as there is no way to reliably tell from the CDR3 sequence which nucleotides originated from V/D/J-segments and which from non-templated additions. We thus removed the nucleotides at the 3’ and 5’ end of the CDR3 that perfectly matched the germline sequence of the annotated TRBV and TRBJ sequence, respectively. The longest match of the remaining sequence with any of the TRBD alleles was taken as a conservative proxy for the D-segment length. We observed a negative relation with incidence in our samples, i.e., sequences shared between samples had on average fewer nucleotides matching a D-segment (Fig. 2C). This indicates that D-deletions may have a positive effect on the abundance in the naive T-cell repertoire.

As described above, there is no method to accurately measure D-segment length of any given TCR*β* rearrangement. We evaluated the performance of our method by applying it to an *in silico* repertoire generated with IGoR [11]. This tool allows to generate TCR*β* sequences with gene choice, deletion and insertion probabilities that are trained on sequence data. The advantage of such generated sequences is that the true D-segment length is known for each recombination scenario. Overall, only 38% of the predictions of our method was correct, which was mainly due to an overestimation of the D-segment length, especially for short D-segments (Fig. 2B). Intuitively these results can be understood because any non-templated insertion will match at least one nucleotide in any of the TRBD alleles. This means that it is very unlikely to observe the complete absence of the D-segment, implying that potential absence of D-segments in TCR*β* sequences is likely overlooked.

To further investigate the role of potential D-segment absence on the abundance of TCR*β* sequences in the naive T-cell repertoires of young individuals, we performed a dual analysis. We started conservatively, by analyzing which rearrangements did not contain any nucleotide encoded by D-segments according to our inference method (remember that the analysis of the *in silico* repertoire shows that the vast majority of such sequences are expected to truly not have D-segment nucleotides (Fig. 2B)). Overall, this feature is relatively rare in our samples (< 0.1%), but much more common among sequences with higher abundance in the naive repertoire (Fig. 2D). The other approach is based on the observation that rearrangements with an inferred D-length of 1 or 2 nucleotides have a > 50% probability of lacking a D-segment (Fig. 2B). Such rearrangements, with 2 or fewer nucleotides matching a D-segment, made up an even larger fraction of the abundant sequences (Fig. 2E). Together, this suggests that there is a substantial fraction of the TCR*β* repertoire of naive T cells that does not contain a D-segment.

### 2.3 TCR*β* sequences without D-segment are abundant, functional, and public

The abundant TCR*β* sequences in the naive repertoire are enriched for having high generation probabilities (Fig. 1A) and also for having no D-segment (Fig. 2D&E). We investigated the relative contribution of these factors in more detail by studying TCR*β* sequences that were abundant (i.e., those that were shared between subsamples). Their generation probabilities were orders of magnitude higher than sequences present in a single subsample (Fig. 3A). The abundant sequences without a D-segment had even more increased median generation probabilities (Fig. 3A). This reveals that the recombination model, of which individual probabilities were trained on large samples of TCR sequencing data, predicts that (almost) complete absence of the D-segment is likely to occur during TCR*β* rearrangement.

**Figure 3:**
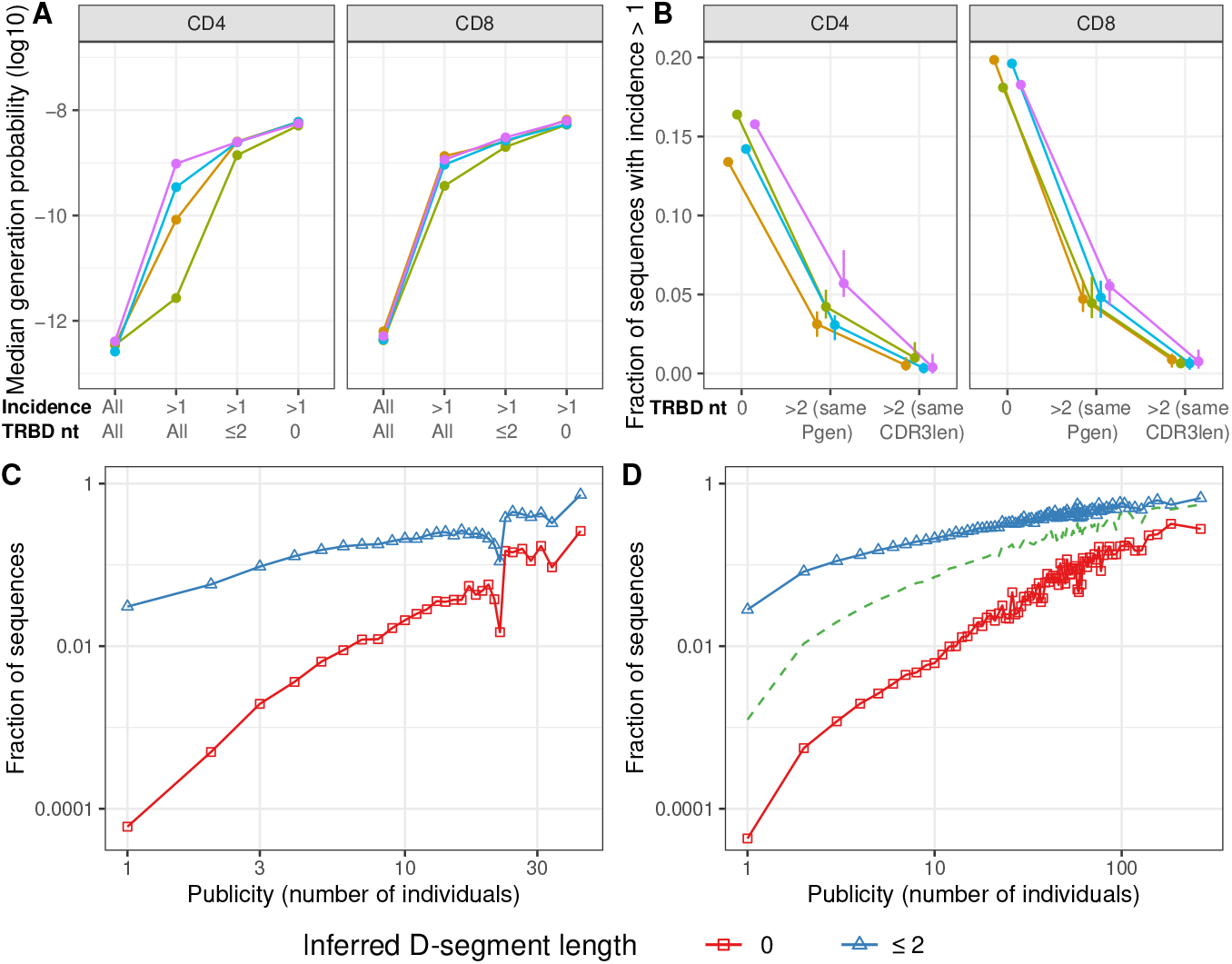
TCR*β* sequences without D-segment are more abundant and shared between individuals. **A.** Median TCR*β* generation probabilities as a function of incidence and inferred D-segment length. **B.** Fraction of sequences occurring in multiple subsamples of naive T-cells. The sequences with an inferred D-segment length of 0 nucleotides are compared with other sequences with an inferred D-segment length of more than 2 nucleotides but identical generation probability or CDR3 length distribution. Closed circles show median of 10 iterations, total range is indicated with vertical bars. **C&D.** The fraction of sequences with an inferred D-segment length of 0 nucleotides (red rectangles) and 2 or fewer nucleotides (blue triangles) as a function of publicity. Publicity values are measured as the number of samples the TCR*β* sequence appears in and are binned such that every data point is based on at least 100 TCR*β* sequences and shown as a weighted average. Data from Britanova *et al.* (C, 73 individuals), and Emerson *et al.* (D, 666 individuals). The green dashed line in D indicates the fraction of reported VJ rearrangements (see Methods).

At the same time, the high generation probability of TCR*β* sequences without D-segment identifies generation probability as a likely confounding factor when assessing the effect of D-segment absence on abundance in the naive repertoire. To discriminate between the effect of both factors, we calculated which fraction of the sequences with an inferred D-segment length of 0 nucleotides was abundant, i.e., shared between subsamples of naive T cells. This fraction was relatively consistent between individuals and between CD4^+^ and CD8^+^ samples (~15% and ~19%, respectively; Fig. 3B). Next, from each set of samples, we selected TCR*β* sequences with more than 2 nucleotides inferred D-segment length, but similar generation probabilities. These sequences, that most likely contain D-segment nucleotides were not nearly as often abundant (Fig. 3B). We performed a similar analysis by taking sequences with identical CDR3 lengths, as absence of the D-segment would lead to shorter CDR3s, which could be another confounding factor. We found that CDR3 length-matched sequences with D-segment were even less often shared between subsamples (Fig. 3B). So, in addition to generation probability and CDR3 length, absence of a D-segment is on its own an important factor affecting the abundance of TCR*β* sequences in the naive repertoire.

The ubiquity of TCR*β* rearrangements in the naive repertoire lacking a D-segment raises the question if such receptors are functional. We therefore assessed their presence in the memory samples from the same dataset and correlated this with incidence among these samples. The fraction of sequences with an inferred D-segment length of 0 or ≤ 2 nucleotides appeared similar between naive and memory samples (Fig. S2A&B), suggesting that absence of the D-segment does not affect the probability of participating in an immune response. The strong relation with incidence that was observed for the naive samples, was however absent for samples of memory T cells. We performed a similar D-segment inference method on the human entries in the VDJdb of reported antigen-specific TCR amino acid sequences [19]. Interestingly, commmon pathogens, like InfluenzaA, EBV, and CMV seem to evoke more responses lacking a D-segment than HIV-1, a more rare pathogen that is typically encountered later in life (Fisher’s exact test, *p* = 0.002, Fig. S2C). Together, these results indicate that TCR*β* sequences without D-segment are not functionally impaired.

The observation that absence of the D-segment causes TCR*β* sequences to be abundant within individuals, predicts that such sequences may also be more often shared between individuals. We tested this by analyzing intra-individual sharing and D-segment lengths of productive TCR*β* sequences from two published TCR*β* datasets of 73 [2] and 666 [5] individuals. In both cohorts, there was a striking relation between publicity and the inferred presence of the D-segment in the CDR3. While our conservative estimate of complete absence of any D-segment was very rare in private sequences (<0.01%), this was the case for about 25% of the most public sequences in both datasets (Fig. 3C&D). We obtained a similar enrichment using the less strict threshold of 2 nucleotides inferred D-segment length (~ 3% versus ~ 70%). This confirms that TCR*β* sequences without a D-segment are not only abundant within the naive repertoire, but also more likely shared between individuals.

### 2.4 TCR*β* sequences without D-segment are generated before birth and maintained until old age

We wondered why especially TCR*β* sequences without D-segment are abundant in the naive repertoire of young individuals. An explanation could be that they were generated prenatally, when clonal competition may be less restrictive. To test this idea, we studied the samples previously described by Carey et al. [3]. They sorted CD8^+^ naive T cells from samples of cord blood from extremely preterm and term neonates and peripheral blood from infants and adults. The cord blood samples contained a much larger fraction of TCR*β* sequences with an inferred D-length of 0 or ≤ 2 nucleotides than the peripheral blood samples (*p* < 0.001; Fig. 4A). Strikingly, the extremely preterm samples contained significantly more sequences without D-segment than the term samples (*p* = 0.016). In contrast, there was no difference between peripheral blood samples of infant and adult individuals. This indicates that generation of sequences without a D-segment is most likely during early fetal development.

**Figure 4:**
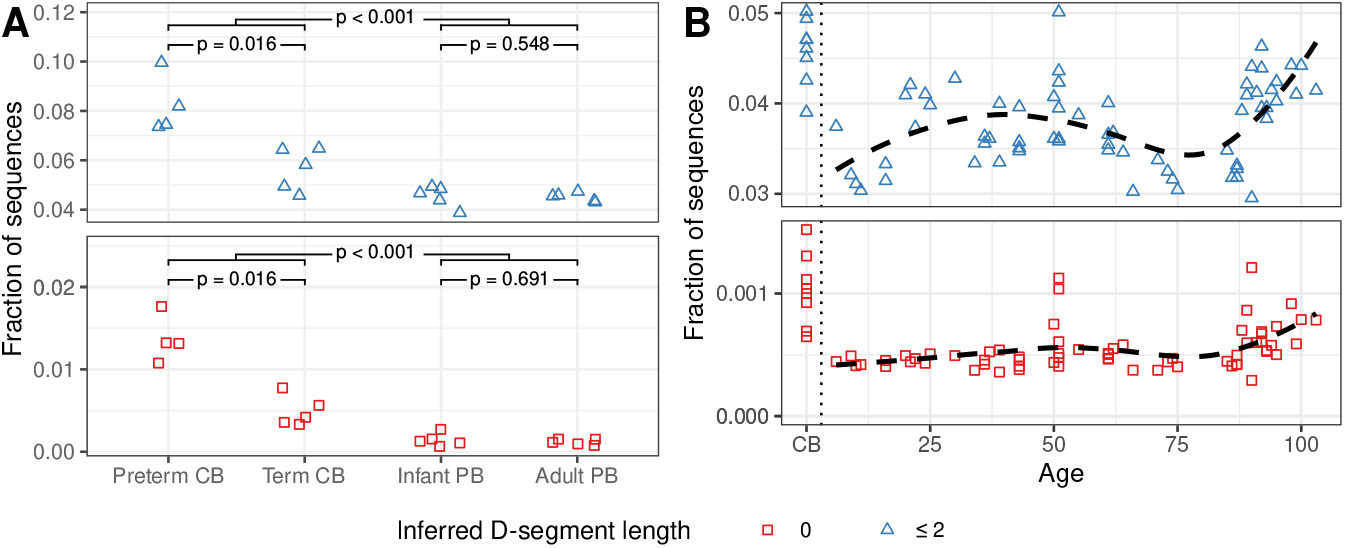
TCR*β*s without D-segment are preferentially generated before birth. **A.** Fraction of TCR*β* sequences with an inferred D-segment length of 0 (red rect1angles) and 2 or fewer nucleotides (blue triangles). Samples of naive CD8^+^ T cells from cord blood (CB) and peripheral blood (PB), as described in Carey *et al.*. The p-values are determined with the Mann-Whitney U test. **B.** D-segment lengths like in A., measured in samples of unsorted T cells, as a function of the individual’s age. Data from Britanova *et al.*. Dashed lines indicate local polynomial regression (loess) on peripheral blood data.

We tested the persistence of such sequences by correlating the fraction of sequences lacking a D-segment with age in the Britanova *et al.* dataset [2], which includes 8 cord blood samples. It should be noted that these samples contained PBMCs that were not sorted to only include naive T cells. In line with the previous results, we found an increased fraction of sequences without D-segment in the cord blood samples. The data from peripheral blood samples indicates that the fraction of sequences with a D-segment length of 0 or ≤ 2 nucleotides stays relatively stable with age, but tends to increase in the elderly of > 80 years. Since there is not much thymic production of new T-cell clones at old age, this likely indicates that TCR*β* sequences without D-segment persist longer than other sequences. Together, these results suggest that TCR*β* sequences without a D-segment are preferentially generated prenatally, dilute before and after birth, but are maintained until very old age.

## 3 Discussion

Here we analyzed TCR*β* sequencing data from naive, memory and unsorted repertoires to identify sequence characteristics that correlate with abundance. We first confirm that abundant TCR*β* sequences in naive T-cell samples of young individuals are characterized by high generation probabilities [17, 23, 14, 4]. Second, we show that the V- and J-gene segments, as well as CDR3 amino acids, that are most common in the overall repertoire, tend to be even more enriched among abundant TCR*β* sequences. These observations indicate TCR*β* sequences being abundant due to repeated production, i.e., reflecting multiple TCR*αβ* combinations that share the TCR*β* sequence and together are more abundant than others [4]. In the aged individuals, these factors correlate with abundance to a lesser extent. This is partly explained by the decreased thymus production in the elderly and indicates some T-cell clones are preferentially selected in the periphery due to other factors.

The most striking observation in our data is the exception to the rule, i.e., glycine, which is due to the absence of the D-segment in many abundant TCR*β* sequences in the naive repertoire of young individuals. Although it is not possible to reliably measure the number of CDR3 nucleotides originating from the D-segment, we used a conservative method and evaluated its performance on an *in silico* repertoire. The D-segment length inference of individual sequences is not very reliable, but our method is conservative and we find convincing evidence for a substantial population of TCR*β* sequences with complete absence of the D-segment in the naive T-cell repertoire. We show that such sequences tend to be much more often abundant in the repertoire and, as a result, more often shared between individuals than other TCR*β* sequences.

From our sequencing data, we cannot infer the recombination scenario by which sequences without D-segment were generated. Following the 12/23 rule, direct V-to-J recombination is possible during TCR*β* rearrangement, although several studies reported this to be rare due to the beyond 12/23 restriction. Still, such a scenario cannot be excluded given the enormous number of TCR*β* rearrangement events during a human lifetime. Alternatively, a large number of deletions at the 3’ and/or 5’ end could remove all D-segment nucleotides. A possible scenario is that N-additions normally ‘protect’ the D-segment against excessive deletion. The TdT enzyme, that is responsible for inserting N-additions during V(D)J-rearrangement, is downregulated during early ontogeny [14], which could make complete deletion of the D-segment more likely, and would explain the large fraction of sequences without D-segment in the cord blood samples from extremely preterm neonates. As a result, complete deletion of the D-segment would become less likely once TdT is activated, causing the rapid dilution of TCR*β* rearrangements without a D-segment even before birth. By this time, the TdT-independently generated clones may have undergone multiple rounds of division, increasing their abundance in the naive repertoire [6], which may be one of the reasons why such rearrangements persist over a human lifetime and even increase in relative frequency in the elderly.

Although our main goal was to describe which sequence characteristics explain abundance in the naive T-cell repertoire, the memory T-cell samples also contain TCR*β* sequences without a D-segment. The abundance in the memory repertoire, however, is not affected by absence of the D-segment. Abundant TCR*β* sequences in the memory compartment likely reflect large clonal expansions rather than the more subtle differences within the naive repertoire. Still, the existence of sequences without D-segment in samples of memory T cells indicates that such rearrangements are functional and participate in immunological responses. Moreover, we find that about 1% of the reported TCR*β* sequences specific for common pathogens does not have any D-matching amino acid in the CDR3 (Fig. S2C). The observation that this percentage is higher for common viral pathogens than for the more rare HIV-1 makes it tempting to speculate about the effect of age at which individuals get exposed to the pathogen. Most people get exposed to common pathogens at young age, when a relatively large fraction of naive T cells originates from prenatally generated clones. Exposure to HIV-1 is much more likely when these sequences are strongly diluted already. If this were the case, it would explain the higher generation probabilities of TCRs specific for common antigens without needing the previously suggested evolution of the recombination machinery towards TCRs specific for common pathogens [20].

Together, our study highlights absence of the D-segment as an important determinant for TCR*β* abundance in the naive T-cell repertoire. Many of them are likely generated long before birth, when TdT is still down-regulated. Such sequences are often shared and maintained until very old age, indicating that the TCR repertoire starts and ends with TCR*β* chains that resemble TCR*α* chains.

## 4 Methods

### 4.1 Data sources

The dataset by Qi et al. [15] was obtained from dbGaP found at https://www.ncbi.nlm.nih.gov/projects/gap/cgi-bin/study.cgi?study_id=phs000787.v1.p1 through dbGaP study accession number PRJNA258304. These data (project “Immunosenescence: Immunity in the Young and Aged”) were provided by Jorg Goronzy on behalf of his collaborators at PAVIR and Stanford University. In this study, five replicates with each 10^6^ cells per aliquot of naive and memory CD4 T cells were collected. For CD8 T cells, 0.25 × 10^6^ T cells were collected per replicate, except for the naive CD8 T cells from young individuals, from which 10^6^ cells per aliquot were analyzed. The TCR*β* repertoire sequence data of 666 indivuduals was downloaded from the Adaptive Biotechnologies website (originally published in Emerson *et al.* [5]). Carey et al. [3] data of cord blood and peripheral blood TCR*β* repertoires were downloaded from the same website. The results presented in Fig. 3C and Fig. 4B are based on data from Britanova *et al.* [2] downloaded from the NCBI SRA archive Bioproject accession PRJNA316572.

### 4.2 Sequence analysis

After pairing reads with Paired-End reAd mergeR (PEAR) [26], the reads from Qi *et al.* were processed using Recover TCR (RTCR) [7]. Each sample of naive or memory cells was split into five subsamples that were sequenced separately. RTCR estimates a per-sample error rate to account for the inevitable inaccuracies that occur during PCR and sequencing errors. To prevent the occurrence of high-incidence reads by the error-correction clustering algorithm, we processed each subsample separately. When running RTCR, we did not perform Unique Molecular Identifier (UMI)-guided error correction. This was because the incorporated UMI sequences were composed of only 4 nucleotides. Thus there are only 256 unique combinations possible, which does not allow for collapsing of PCR duplicates into reliable consensus sequences. As we did not use within-sample read counts, but only used the incidence in multiple samples as a measure for abundance, our results should not be influenced much by uneven PCR amplification. After error-correction, reads were filtered following default RTCR settings, i.e., if V and J were in-frame and the CDR3 did not contain in-frame stop codons or ambiguous bases. TCR*β* sequences were defined by the combination of CDR3 nucleotide sequence, TRBV gene and TRBJ gene.

The data by Britanova *et al.* was processed with RTCR using the barcode files given at https://github.com/mikessh/aging-study. First, UMIs were extracted in forward and reverse reads using the Checkout algorithm of RTCR. UMI-guided consensus sequences were generated using the umi_group_ec algorithm, which were processed with the main pipeline of RTCR using default settings.

The data from Emerson *et al.* was already processed. TCR*β* sequences were extracted from the column “rearrangement”, V and J genes from the columns “v_gene” and “j_gene”, respectively. Out-of-frame sequences, those with an unresolved V or J gene or containing an in-frame stop codon, were filtered out and not used for analysis. Publicity was measured as the number of samples that contained the combination of CDR3 nucleotide sequence, V gene and J gene. The fraction of VJ rearrangements (green dashed line in Fig. 3D) is based on the column “rearrangement_type”.

For analysis of the Carey *et al.* data, TCR*β* CDR3 nucleotide sequences were taken from the column “cdr3_rearrangement”, V and J genes from the columns “v_gene” and “j_gene”, respectively. Out-of-frame sequences, those with an unresolved V or J gene or containing an in-frame stop codon, were filtered out and not used for analysis.

### 4.3 Inference of D-segment length

Many different recombination scenarios can lead to the exact same sequence, e.g., after deletion of nucleotides they can be added again as N-insertion. As these differences are not visible in the sequence, it is impossible to tell which nucleotides are encoded by V, D or J segment, and which by N-additions. To still estimate the number of nucleotides originating from the D-segment, we used a conservative approach. We start by taking the CDR3 nucleotide sequence and matching the nucleotides at the 5’ end to the germline sequence of the identified V gene segment. The first mismatch position is assumed to be the end of the V-segment, although technically this mismatch could also occur due to e.g. a sequencing error. The same procedure is followed by matching the 3’ end of the remaining sequence to the identified J gene segment. Any remaining nucleotides could be a mixture of a D gene segment, N-additions and P-additions. We inferred the length of the D-segment by taking the longest exact match of any of the three germline TRBD allele sequences (as listed in IMGT [8]: TRBD1*01:GGGACAGGGGGC; TRBD2*01:GGGACTAGCGGGGGGG; TRBD2*02:GGGACTAGCGGGAGGG) with the inter-V-J sequence.

To evaluate the performance of the D-segment length inference method, we generated 10^7^ TCR*β* sequences using the default generation model of IGoR without sequencing errors. We randomly selected 10^6^ sequences that were productive, i.e. in-frame and not containing a stop codon, and which V- and J-segments were in the RTCR germline reference set (e.g., excluding pseudogenes). For each sequence, we inferred the D-segment length as described above. We compared this to the true D-segment length, by subtracting d_5_del and d_3_del (if positive) from the total length of the selected D-segment.

---

**Figure S1:**
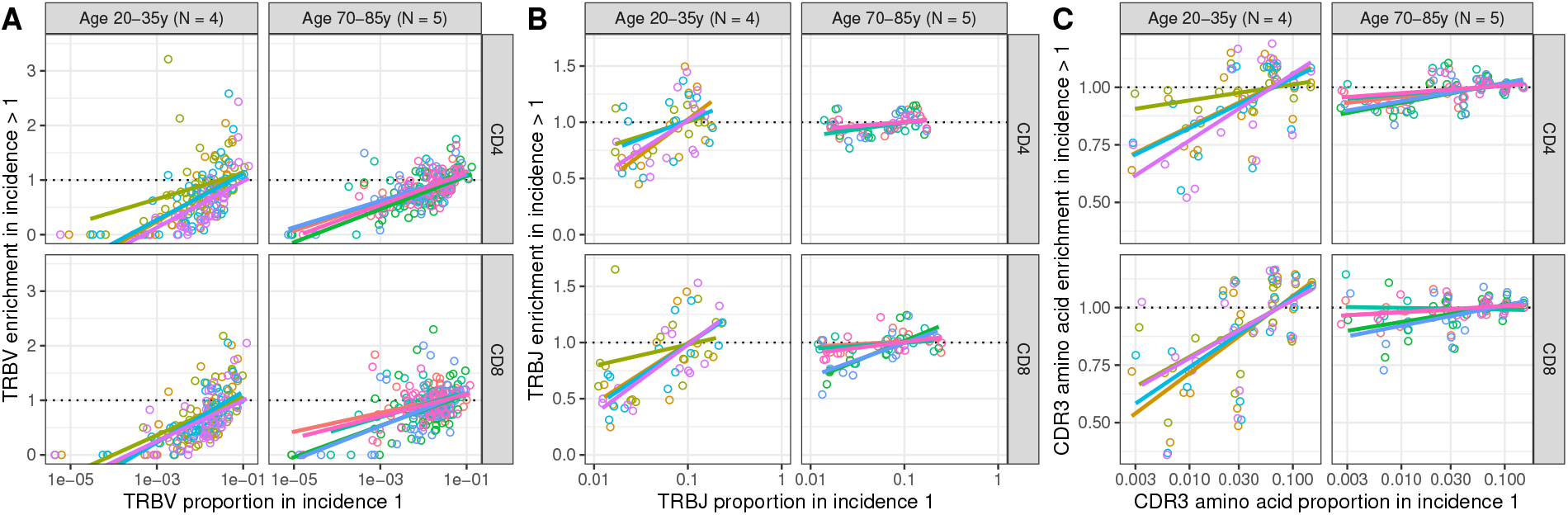
Common V/J segments and CDR3 amino acids are enriched in abundant TCR*β* sequences. Enrichment values like in Fig. 1 as a function of their prop1ortion among sequences with incidence 1 (log-scaled). Each color represents a different individual, matching the colors in the main text figures. The positive slopes of the linear regression lines indicate that common segments and amino acids tend to be enriched in abundant sequences.

**Figure S2:**
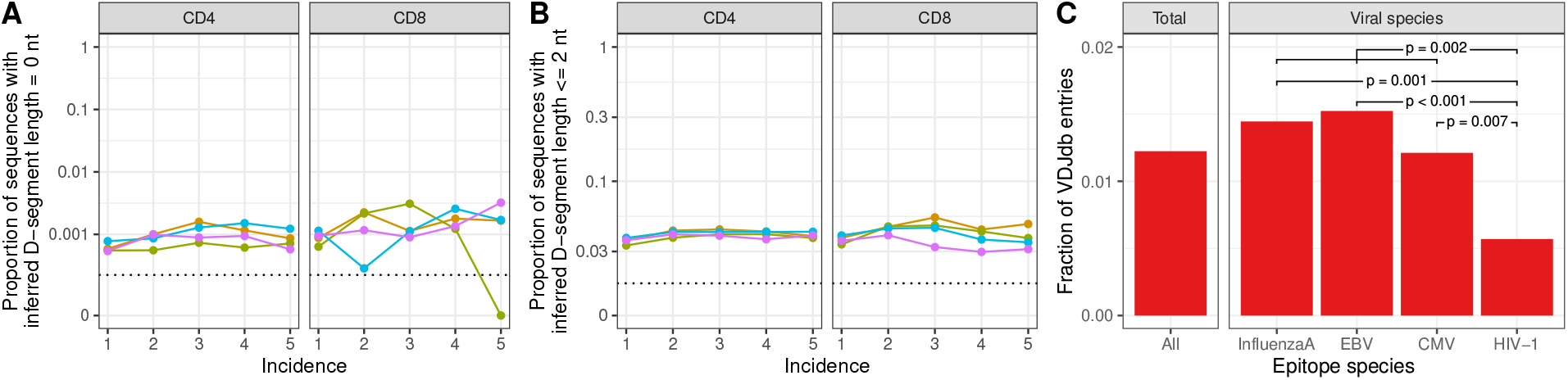
TCR*β* sequences without D-segment occur in memory and antigen-specific T cells. **A&B.** Fraction of sequences with an inferred D-segment length1 of 0 (A) and 2 or less (B) nucleotides as a function of incidence in memory samples. The vertical axes have logarithmic scales with 0 added at the bottom of the axis. **C.** Inference of D-segment length on CDR3 amino acid sequences in the VDJdb, retrieved on 8 January 2021 [19]. Like for nucleotide sequences (Methods), we assigned matching CDR3 amino acids to the translated germline V and J sequences. In the remaining amino acids, we used the maximum match with any of the reading frames of translated TRBD alleles (Fig. 2A) as a proxy for the number of D-encoded amino acids in the CDR3. Shown are the fractions of unique sequences having 0 D-encoded amino acids among all human TCR*β* records (left), and those specific for the viral epitope species with at least 1000 unique V-CDR3-J combinations (right). P-values are determined with Fisher’s exact test.

